## Supplementary material for "Balance of Mechanical Forces Drives Endothelial Gap Formation and May Facilitate Cancer and Immune-Cell Extravasation"

#### Supporting Information, Text

#### Detailed derivation of the model

The endothelial monolayer is modeled through a number of cells in two dimensions that are connected through cell-cell adhesions. Each cell contains a number of radial contractile actin stress fibers, modeled by viscoelastic springs. Then, viscoelastic springs also represent the combined cell membrane with the adjacent cortex. For simplicity, in the paper we will refer to these elements as membrane elements. Therefore,the whole cell is discretized into nodes, which represent the fundamental degrees of freedom of the resulting network of stress fibers, membrane elements and cell-cell junctions. The cell membrane is discretized into $n_{nodes}$ nodes with a spacing, $l_n$ between them. The example of a hexagonal geometry is shown in Fig. S1, where all radial stress fibers are connected in the center of the cell. The actual model is independent of cellular geometry. In fact, cell geometry is dynamically changing, as observed in experiments (Movie S2), and that is reflected in the model simulations. Both membrane and stress fibers are considered as viscoelastic elements and are approximated by Kelvin-Voigt structures, in line with previous models [28]. We assume that inertial forces have no significant impact on the system, as typical on the cellular scale [54], and the forces exhibit inherent randomness due to fluctuations in molecular activities in each cell and due to variability in growth factors or neighboring cells that inevitably affects cell mechanics. The dynamics of each node is then described by a Langevin equation (where inertia effects have been neglected), similar to the one that appeared in other cell mechanical models (see e.g. [54]):

$$\frac{d\mathbf{r}_i}{dt} = \frac{1}{\xi}\mathbf{F}_i \tag{S1}$$

Here, $\mathbf{r}_i$ corresponds to the current position of each membrane node and cell center $i$, $\xi$ is the medium drag coefficient, and $\mathbf{F}_i$ represents the sum of all forces due to active, random contractions or through passive mechanical interactions between node $i$ its neighboring nodes:

$$\mathbf{F}_i = \mathbf{F}_i^{sf} + \mathbf{F}_i^{memb} + \mathbf{F}_i^{adh} + \mathbf{F}_i^{rep} + \mathbf{F}_i^{gen}, \tag{S2}$$

where, $\mathbf{F}_i^{sf}$ is the force due to radial stress fibers, $\mathbf{F}_i^{memb}$ is the force due to the membrane and cortex, $\mathbf{F}_i^{rep}$ is the repulsion force due to contact between different cells, $\mathbf{F}_i^{gen}$ is the cell generated force due to contractions or protrusions of the stress fiber or membrane elements, respectively. $\mathbf{F}_i^{adh}$ is the force originating through cell-cell adhesions, represented by VE-cadherin in our stochastic adhesion model described below.

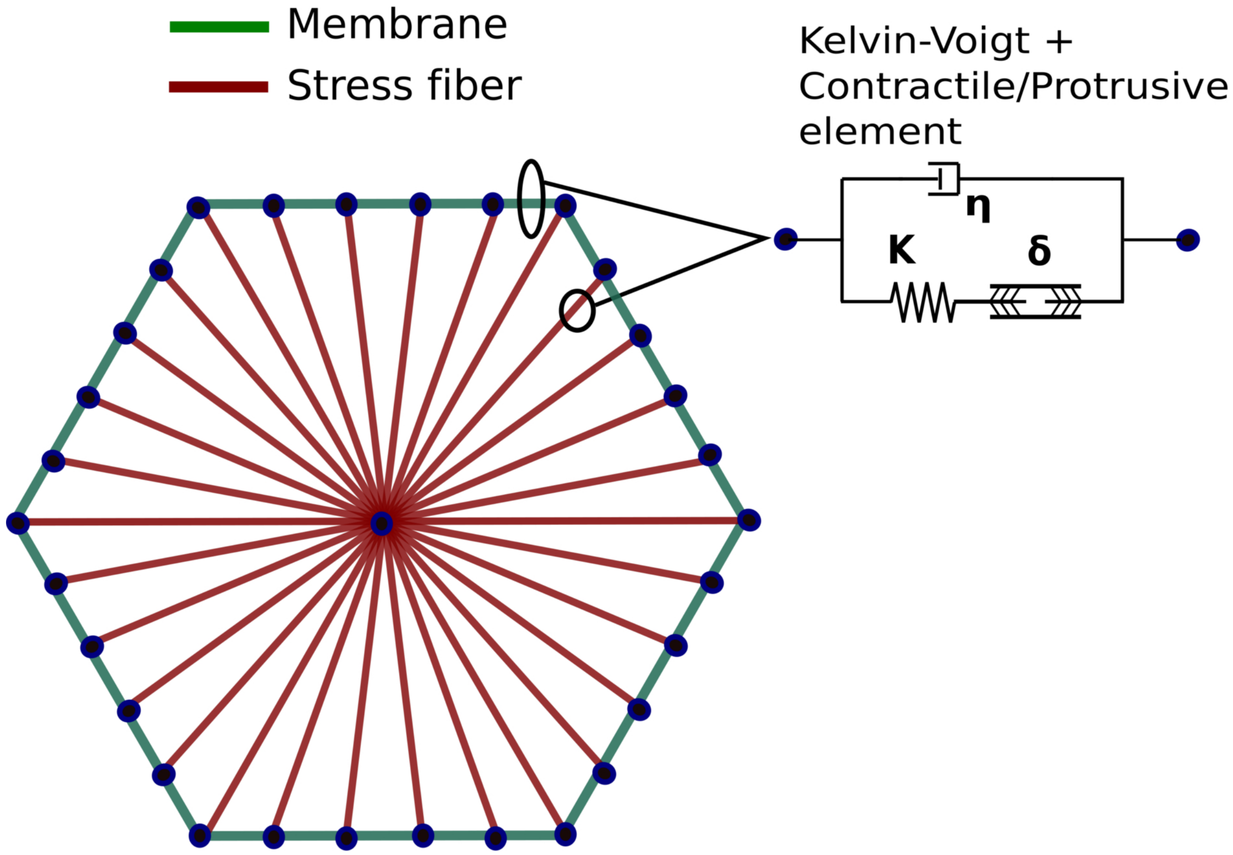


Figure S1: **Mechanical model of a single cell.** The cell is presented in an initially hexagonal form, divided into a discrete number of membrane points. Physically, our membrane elements connecting the nodes represent the combined lipid bilayer with the actin cortex. Moreover, the nodes are connected to the center by stress fiber structure. Both of them are described by Kelvin-Voigt models with a contractile/protrusive element, but both have different parameters.

#### Model of Passive Intracellular Mechanics

We now describe all mechanical properties modeled within a single cell. As described before, both radial stress fibers and tangential membrane/cortex segments are modeled with Kelvin-Voigt elements (S1). For

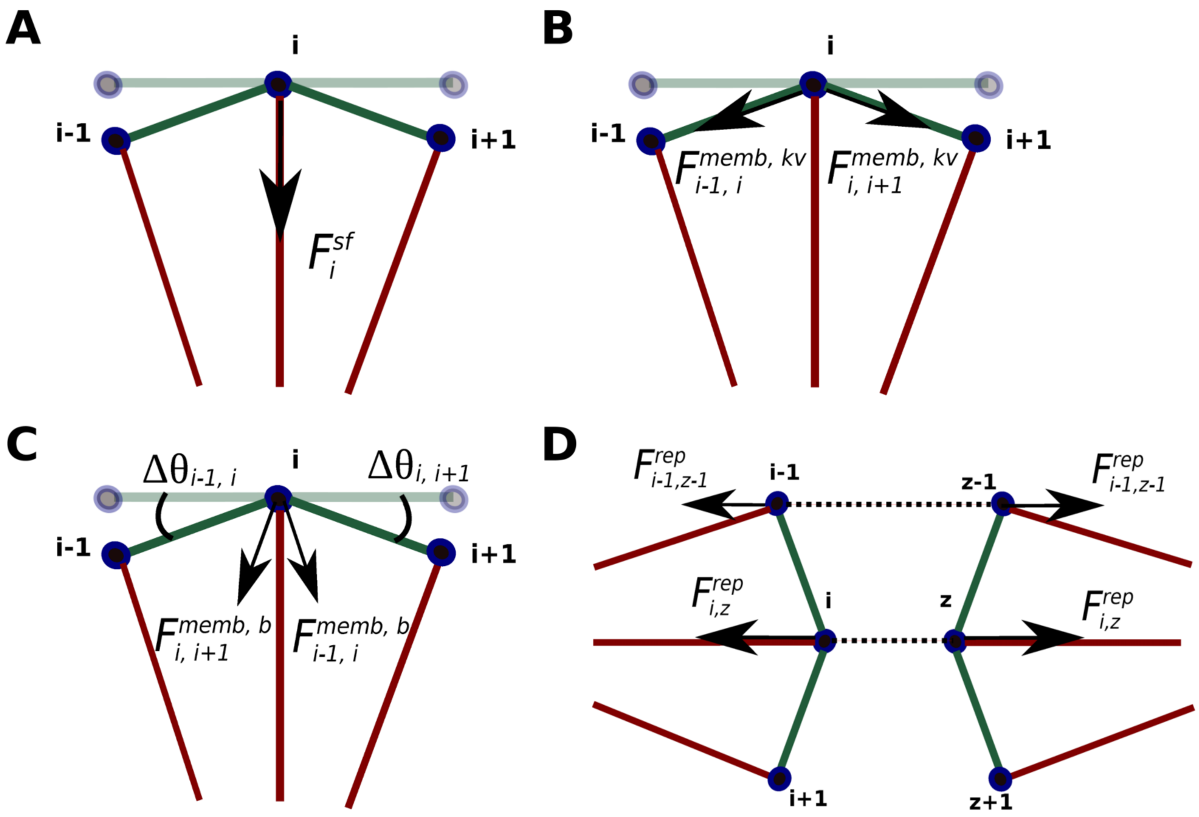


Figure S2: **Contribution of different passive intracellular forces.** (A) Force due to stress fiber deformations. (B) Force due to membrane in-plane deformation. (C) Force due to membrane bending stiffness. (D) Force due to repulsion between membrane points of different cells.

simplicity, we consider hexagonal cells as a starting point, even though the actual modeling framework is independent of the cell geometry. Stress fibers connect a node in the center of the cell with a node on the membrane. Membrane and center nodes are treated in the same way. Additional to the Kelvin-Voigt force arising due to deformations in the direction of two membrane points $\mathbf{F}_i^{memb,kv}$ (Fig. S2B), the membrane/cortex exhibits bending stiffness resulting in forces due to deformations perpendicular to the membrane, $\mathbf{F}_i^{memb,b}$ (Fig S2 C). The total force on a membrane node due to deformations of neighboring membrane nodes is thus given by:

$$\mathbf{F}_i^{memb} = \mathbf{F}_{i-1,i}^{memb,kv} + \mathbf{F}_{i+1,i}^{memb,kv} + \mathbf{F}_{i-1,i}^{memb,b} + \mathbf{F}_{i+1,i}^{memb,b} \tag{S3}$$

$$|\mathbf{F}_{i-1,i}^{memb,b}| = \frac{K_{bend}}{d_{i-1,i}} \cdot (\beta_{i-1,i} - \beta_{i-1,i}^0), \tag{S4}$$

where $i-1$ and $i+1$ are, without loss of generality, neighboring points of $i$, $\beta_{i-1,i}$ is the angle denoting deviations from the balance position which is $\beta_{i-1,i}^0$ (see Fig. S2C) and $d_{i-1,i}$ is the distance between the two neighboring points. For the forces derived from the Kelvin-Voigt structures, $\mathbf{F}_i^{memb,kv}$ and $\mathbf{F}_i^{sf,kv}$, the direction corresponds to the vector $\vec{\mathbf{k}}$ formed by the corresponding nodes as indicated in the sub-indices ($i-1$ or $i+1$, $i$). For the bending stiffness component ($\mathbf{F}_i^{memb,b}$), the direction of the force is perpendicular to $\vec{\mathbf{k}}$ (unit vector of the membrane segment), and $K_{bend}$ is the rotational spring constant used to approximate membrane bending effects.

Kelvin-Voigt structures consist of a parallel arrangement of an elastic spring and a viscous damper, with the force

$$F(t) = EA\varepsilon(t) + \frac{d\varepsilon(t)}{dt}A\eta. \tag{S5}$$

Here, $E$ is the elastic modulus of the spring, $A$ is the area of the cross-section, and $\eta$ is the viscosity of the material (membrane or stress fiber, respectively). Forces are implemented for both stress fibers and membrane structures as follows:

$$\mathbf{F}_i^{sf} = [K_{sf}(l_n - l_n^0) - \eta_{sf}v_n] \cdot \vec{\mathbf{k}}, \tag{S6}$$

$$\mathbf{F}_i^{memb,kv} = [K_{memb}(l_n - l_n^0) - \eta_{memb}v_n] \cdot \vec{\mathbf{k}}, \tag{S7}$$

where $K_{sf}$ and $K_{memb}$ are the stiffness and $\eta_{sf}$ and $\eta_{memb}$ are the drag coefficients of stress fibers and membrane, respectively. $n$ denotes the bar in the network that connects the points $i$ and $i-1$, and could correspond to a stress fiber or a membrane segment. $l_n$ is the current length of the element, and $l_n^0$ is the rest length. $v_n$ corresponds to the velocity at which the element is varying its length, and $\vec{\mathbf{k}}$ is the unit vector in the direction of the element.

#### Model of cell-cell junctions

Endothelial cells are mechanically coupled to neighboring cells through cell-cell adhesions. VE-Cadherin is the major protein in endothelial cell adherens junctions and is known to cluster on the membrane [23]. It is a homophilic protein binding to other VE-cadherins on neighboring cells, and also links cell-cell adhesions to the cytoskeleton [30, 22]. Cell-cell adhesions in endothelial cells are very complex, and also include tight junctions. In our model, the precise molecular composition and regulation of the junctions is not relevant. However, it is important to note that several molecular bonds in adhesion complexes are force-sensitive. For instance, a force-dependence of VE-cadherin/VE-cadherin bonds at high forces was known for a long time [30], and the same paper showed that these bonds have a longer lifetime than the related E-cadherin or N-cadherin bonds present in other cell types. More recent evidence showed that cadherin bonds may also exhibit a catch-bond nature [55], causing the bond-lifetime to initially increase with force. Likewise, the bonds connecting cadherins to the cytoskeleton, for instance through $\alpha$-catenin, were recently found to be described by a catch bond [56].

Our model is thus designed to capture the force dependence of the cell-cell adhesions. Each discretized membrane point can act as a local adhesion cluster that may bind to a membrane point on adjacent cells. If binding occurs, the cells are physically connected through a linear spring with force

$$|\mathbf{F}_i^{adh}| = |\mathbf{F}_z^{adh}| = K_{adh,p} \cdot (d_{i,z} - L_{adh}^0). \tag{S8}$$

Here, $d_{i,z}$ is the distance between the node $i$ on the cell under consideration, and $z$ denotes the node on the adjacent cell. $K_{adh}$ is the stiffness constant of the adhesion complex and $L_{adh}^0$ is the adhesion equilibrium length. The direction of the force corresponds to the vector formed by the two points of the adhesion, $i$ and $z$. From now on, we refer to the adhesion complex that binds points $i$ and $z$ with the subindex $p$.

The probability of binding of two membrane points is determined by a rate that depends on the distance between these two points:

$$k_{bind,p} = \begin{cases} k_{on}^0 \cdot \rho_{adh} \cdot (1 - \frac{d_{i,z}}{L_{bind}^{limit}}) & \text{if } L_{bind}^{limit} \leq 0, \\ \\ 0 & \text{if } L_{bind}^{limit} > 0, \end{cases} \tag{S9}$$

Here, $k_{on}^0$ it is a binding rate constant and $L_{bind}^{limit}$ is the maximal distance at which two membrane points of neighboring points could bind, and $\rho_{adh}$ is the density of adhesion molecules available for binding.

To describe the experimentally observed effect that adhesion complexes may strengthen due to clustering of molecules such as VE-cadherin and recruitment of molecules such as talin or vinculin, which themselves have force-dependent binding rates that may lead to positive feedback loops, [23], our model includes a force dependence of the adhesion complex density and consequently its mechanical properties. Indeed, it was shown that forces play an active role during the strengthening of cell-cell adhesions [23, 20]. Our model thus incorporates a mechanism to reinforce cell-cell junctions in response to forces. This is done by describing the bond density $n_{reinf}$ through a stochastic, force dependent model. For simplicity, and to effectively describe the important case of $n_{reinf} = 0$ that corresponds to complete rupture of the adhesion complex under consideration, we employ a discrete model where $n_{reinf}$ takes on discrete values between 0 and $n_{reinf}^{max}$. $n_{reinf}^{max}$ corresponds to maximal saturation of the adhesion complex, i.e. maximal binding strength. Each adhesion complex may thus act as a molecular clutch, unbinding for zero density, and fully engaging at $n_{reinf}^{max}$. The resulting stiffness of the spring then changes in a linear way:

$$K_{adh,p} = n_{reinf,p} \cdot K_{adh}^0 \tag{S10}$$

Here, $K^0_{adh}$ is the stiffness per unit bond density.

Once a connection between two neighboring cells is formed, (following eq. S9), the bond density of the adhesion complex can vary stochastically following a force dependent law where the reinforcement rate is given by:

$$k_{reinf,p} = \begin{cases} k^0_{reinf} \cdot \rho_{adh} \cdot (1 - (\lambda_{reinf} - F^{adh}_p)/\lambda_{reinf}) = \\ \\ k^0_{reinf} \cdot \rho_{adh} \cdot F^{adh}_p/\lambda_{reinf} \quad \text{if } F^{adh}_p \leq F^{limit}_{reinf}, \\ \\ 0 \quad \text{if } F^{adh}_p > F^{limit}_{reinf}. \end{cases} \tag{S11}$$

Here, $k^0_{reinf}$ is the binding rate constant for the reinforcement and $\lambda_{reinf}$ is a force constant for shifting the reinforcement curve. Furthermore, $F^{limit}_{reinf}$ is a threshold above which we stop applying the reinforcement. This threshold is set to avoid numerical instabilities due to very large binding rates. It has no physical consequences for the model behavior as long as it is numerically set to much larger values than the force corresponding to the peak lifetime of the catch bond, see Eq. (S12) and Fig. S12 below. This is because for such high forces, the cell-cell adhesion clusters are already certain to have unbound due to the catch-bond nature we are now discussing.

Unbinding of single bonds is modeled as a catch bond law [57]:

$$k_{ub,p} = k^0_c \cdot exp(\theta_c - \theta) + k^0_s \cdot exp(\theta - \theta_s), \tag{S12}$$

where $\theta = |F^{adh}_p|/F^0$ and $\theta_c$, $\theta_s$ are the parameters of the catch and slip bond regimes respectively. $F^0$ is used to normalize the force, $|F^{adh}_p|$ is the modulus of the current force on the specific bond and $k^0_c$ and $k^0_s$ are the unbinding rate for coefficient for the catch and slip curve respectively.

Since the mechanics of our monolayer is described through connected springs, where the dynamics is exclusively calculated through the forces on the nodes, such springs could hypothetically overlap. A repulsion force on membrane nodes is thus included to prevent different cells from overlapping. This force occurs when two membrane points of two different cells are within a certain small distance range ($L_{rep}$). The magnitude of this force grows with the distance between two membrane points, $i$ and $z$, of the two adjacent cells(Fig. S2D):

$$\mathbf{F}^{rep}_{i,z} = K_{rep} \cdot (L_{rep} - d_{i,z}) \cdot \vec{\mathbf{j}} \tag{S13}$$

Here, $K_{rep}$ is a constant parameter, $d_{i,z}$ is the distance between the two membrane points of different cell, and the direction of the force is obtained as in Fig. S2D. $L_{rep}$ is the maximum distance at which repulsion is applied and $\vec{\mathbf{j}}$ is the unit vector in the opposite direction to the straight line that binds both points ($i$ and $z$).

#### Cell-generated forces

Forces are generated within the cell due to motor activity and cytoskeletal remodeling. Myosin generated forces ($\mathbf{F}_i^{myo}$) act on both the stress fibers and the cortex membrane in a contractile manner, and protrusive forces ($\mathbf{F}_i^{prot}$) generated by actin polymerization may lead to forces directed towards the outside of a cell:

$$\mathbf{F}_i^{gen} = \mathbf{F}_i^{myo} + \mathbf{F}_i^{prot} \tag{S14}$$

Myosin forces are the result of the combination of two types of forces. The first one, which is generated by the myosin activity in the stress fibers, thus typically results in radial forces. The second one are tangential forces that occur due to contractions of the cortical actin filaments and are directed parallel to the membrane. Both forces have a magnitude of $F_{Radial}^{max}$ and $F_{Cortex}^{max}$ for each stress fiber and membrane segment respectively. Also, both type of forces are not homogeneously distributed throughout all the stress fibers and membrane segments of the cell and are not acting during the whole simulation time. The spatial distribution of the forces is controlled by $n_{Radial}^{Force}$ and $n_{Cortex}^{Force}$, which represents the number of consecutive stress fibers or membrane segments respectively that have the same force. Each one of this set of segments has a probability for activating the force of $p_{Radial}^{Force}$ and $p_{Cortex}^{Force}$. Depending on this probability, the force for each segment of the different sets is either $F_{Radial}^{max}$, or a baseline contraction, which we simulate to be a random number between $[0,\ 0.1 \cdot F_{Radial}^{max}]$ for the radial force case. For the cortex force, depending on the outcome of our Monte-Carlo simulation, the magnitude of the force is either $F_{Cortex}^{max}$ or zero. This probability is calculated for each of the stress fiber and membrane sets at every given time interval indicated by $t_{Radial}^{Force}$ and $t_{Cortex}^{Force}$. These values indicate the time steps when the force activations are recalculated in the Monte-Carlo simulation (see Fig. S3A,B), and they are much larger than the overall time step used to simulate the whole system (Table S1). If, following the previous explanation, there is a change in the force due to a random recalculation of the forces for either stress fiber or membrane segment, the force does not change abruptly in one time step. Instead, the force magnitude changes linearly in time from the previous value to the new one in a given total time, $t_{Transition}^{Force}$. This is to mimic the behavior of cells while external conditions remain approximately constant, so no rapid changes in the mechanics within each cell occur (compare for Movie S2). In this way, cell forces are not homogeneously distributed in time and space, but also do not change abruptly in the absence of external stimuli. External stimuli, for instance, vasoactive agents like thrombin increase intracellular levels of $Ca^{2+}$ and lead to myosin activation [15]. This induces changes in traction forces, leading to heterogeneous force distribution which causes the formation of inter-cellular gaps.

Protrusive forces are caused due to actin polymerization at the edges of a cell. For simplicity, protrusive forces are modeled in the same way as contractile forces, and are only distinguished from contractile forces due to the direction of force and the characteristic parameters. Thus, they act typically in the opposite direction of radial contractile forces (i.e. outward of the cell) and with their own parameters characterizing the typical magnitudes ($n_{Prot}^{Force}$, $p_{Prot}^{Force}$ and $t_{Prot}^{Force}$) (see Fig. S3C).

#### Actin remodeling

The protrusions due to actin polymerisation, as described above, may change the length of stress fibers. Likewise, shrinkage of existing fibers may occur due to depolymerization, or due to severing or buckling and subsequent breakage of fibers [58]. Other mechanisms leading to changes of the rest length of stress fibers include the addition of sarcomeric units in the middle of stress fibers in response to tension [59]. Our model effectively incorporates the dynamical remodeling of stress fibers due to adaptation to the applied forces. For simplicity, we do not consider total depolymerization of a fiber or de novo polymerization of new fibers in response to nucleation. Moreover, we assume that the total amount of F-actin is conserved in a given simulation, i.e. the G-actin available after depolymerization is assumed to quickly polymerize in other fibers.

We describe the remodeling of the stress fibers through a change in the rest length of the spring in the Kelvin-Voigt element. This way, we do not explicitly take into account the precise origin of the change in rest length (e.g. whether it is due to actin polymerization, depolymerization or inclusion of sarcomeric units). Stress fibers dynamically remodel by adapting their rest length to their current length at a certain velocity:

$$\dot{L_s^0} = v_s^{remodel} = K_{remodel} \cdot (L_s - L_s^0) \tag{S15}$$

Here, $s$ is the index of the stress fiber, $L_s$ is the current length of the stress fiber, $L_s^0$ is the current balance rest length of the stress fiber and $K_{remodel}$ is a constant describing the rate of length adaptation.

Since we assume total F-actin conservation, this means that under constant cross-sectional area the total balance rest length of the stress fibers is constant:

$$\sum_{s=1}^{S} L_s^0 = const \tag{S16}$$

Here, $S$ is the total number stress fibers in a cell. For simplicity, we ignore the spatial variations of actin regulators and assume each stress fiber has a similar amount of free barbed ends that polymerize. In order

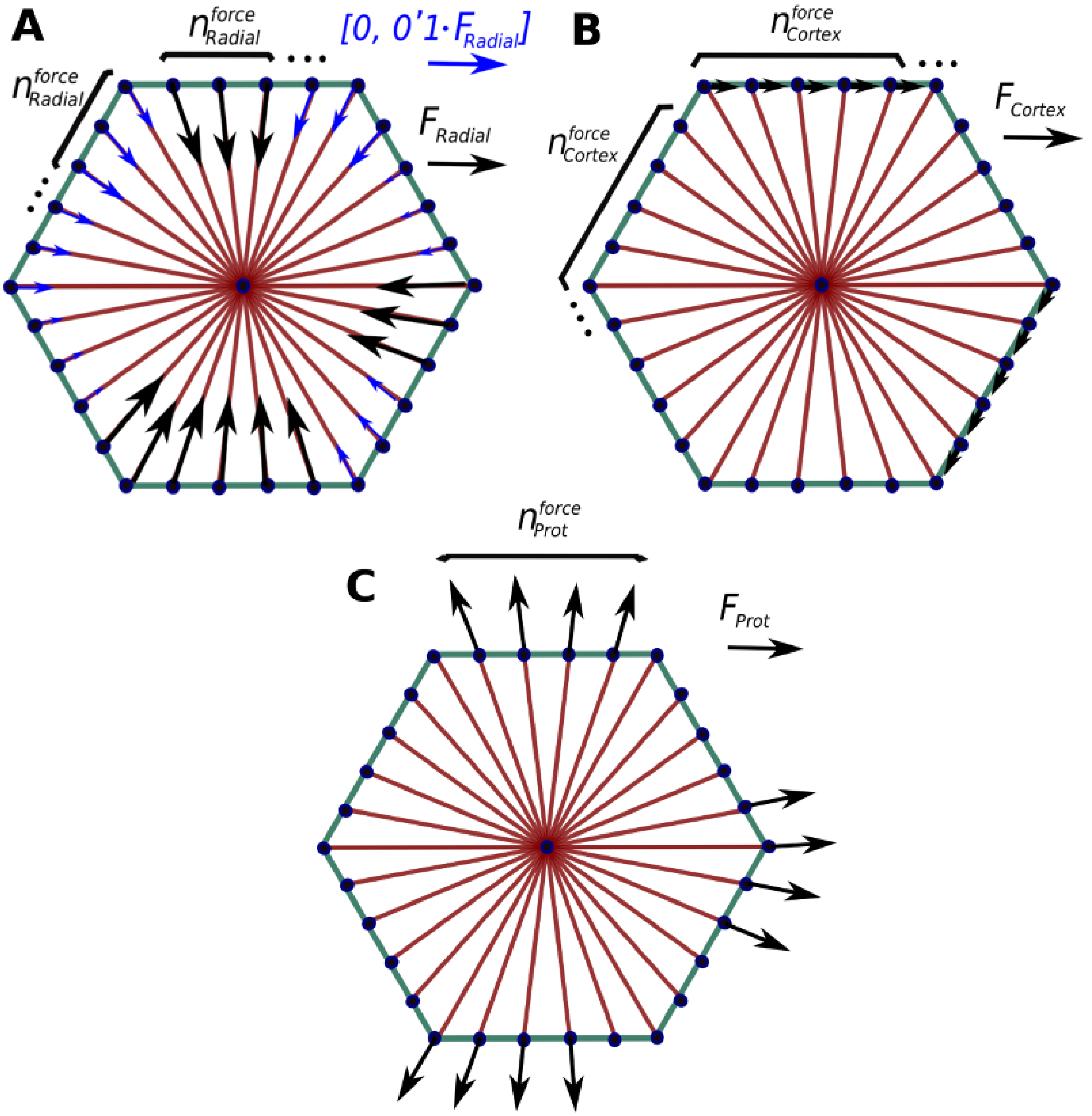


Figure S3: **Cell generated forces.** (A) and (B) Correspond to myosin forces: Radial force and Cortex force respectively. (C) Protrusive forces.

to satisfy Eq. (S16), the rest length of all the stress fibers in a cell is thus modified in the same way (see Fig. S4).

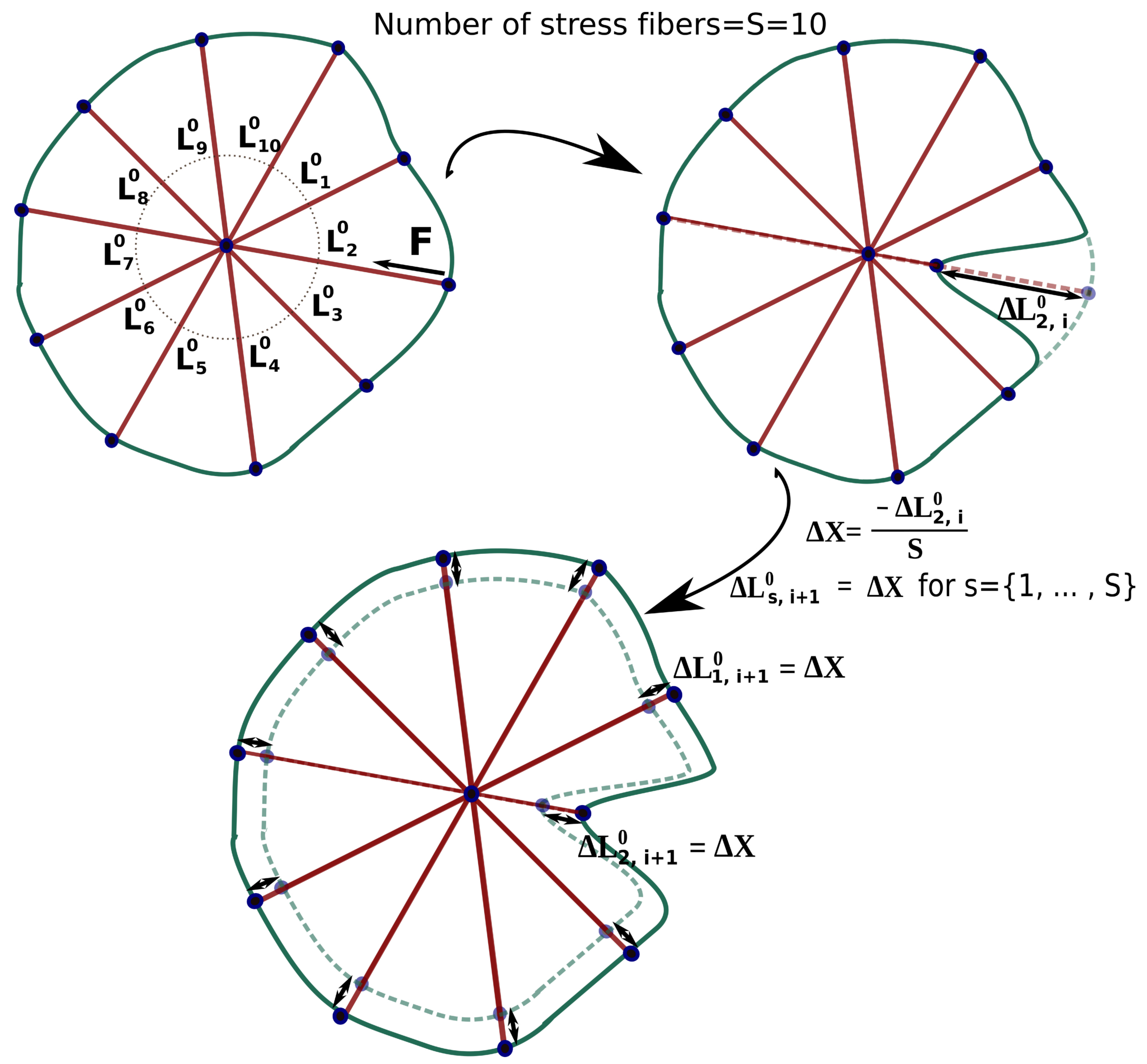


Figure S4: **Stress fiber remodeling.** Due to myosin contractility, a change in the rest length of the stress fiber occurs accordingly to Eq. (S15). This change in rest length is compensated by all the stress fibers in a proportional way. Note that only the rest lengths and not the current length of a stress fiber is modified.

### Implementation of the model and simulations

#### Initial and boundary conditions

Fig. S5 shows an example of a cell monolayer, composed of initially hexagonal cells. Here, the monolayer is initialized such that all neighboring adhesions are bound. Membrane points at the edge of the monolayer are encastred, so that the total domain of the monolayer is fixed. This is to mimic our experimental conditions where the cells are placed in fixed devices. The aim of the simulations is to study the mechanisms behind gap formation. To avoid effects due to the boundary conditions, our gap quantification is performed for the cell in the center of the monolayer only. The initial conditions are such that all spring elements are at balance. When the simulation starts, myosin forces and protrusive forces activate and disturb the balance.

#### Quantification of gap formation

Gaps are formed between two or more cells as a consequence of the adhesion bond rupture. In the model, as described above, two cells are connected at two adjacent nodes through adhesion complexes that are characterized through a (for simplicity assumed discrete) number of bonds $n_{reinf}$. If $n_{reinf} = 0$, the adhesion complexes of the adjacent cells unbind. However, the unbinding of a single pair of adhesion complexes on adjacent cells does not necessarily imply that the endothelial barrier is breached at that location, as this requires a sufficiently high number of close-by adhesion complexes to rupture. We thus quantify the breached area in between two or more cells that resulted from ruptured bonds and only quantify the rupture events as the formation of a proper gap if the area exceeds a threshold area $A_{GAP,F}$. Likewise, when the gap area drops below a threshold $A_{GAP,C}$, we consider that gap to be closed. $A_{GAP,C}$ is typically chosen to be slightly smaller than $A_{GAP,F}$, since otherwise randomness in the simulation may lead to fluctuations around this detection threshold and thus incorrectly predict gaps to form and close constantly. We differentiate between gaps that are formed at a two cell border and gaps that are formed in the vertex of the cells touching three or more cells. A typical gap and how it is quantified is shown in Fig. S6.

#### Implementation

The model has been implemented in a custom-made C++ program. We used the Eigen3 library for operation with the vectors and matrices used to solve the equations. For the rest of the program we have used standard C++ libraries. The code is available on Github, https://github.com/Escribs/Endothelial-monlayer.

#### Simulations

In each simulation, we first set up the initial and boundary condition and then calculate a loop consisting of repeatedly executing the following major steps:

1. Binding and reinforcement of adhesions: First we analyze the position of membrane points of different cells and calculate the probabilities for binding. We perform a Monte-Carlo simulation to see if a new union is formed. We also analyze the reinforcement of adhesions that are already bound.

2. Node displacement: We analyze the force balance in each node $i$ of the monolayer and calculate the movement of each node with the Langevin equation (S1). The forces considered are the ones outlined

in Eq. (S2). To avoid instabilities, we set a maximum node displacement. If the node displacement exceeds this threshold, the time step is dynamically reduced when integrating the Langevin equation; a process we perform iteratively. In Fig. S15, we show that the default time step we fixed is sufficiently low so that simulation results are not significantly affected by this time step.

3. Actin Remodeling: Based on the new positions of the nodes, we simulate the remodeling of actin.

4. Unbinding of adhesion bonds: We update the forces on adhesions after the displacement of nodes, and preform a Monte-Carlo simulation to check if bonds unbind according to Eq. (S12).

5. Gap formation: Finally we quantify the formation of new and closure of existing gaps and the size of all currently existing gaps.

#### Parameter Justification

The parameters of the reference case are summarized in Table S1. Parameters for the unbinding law are adjusted to match data from [30], where distributions of forces were shown to be required to break single VE-cadherin/VE-cadherin bonds in HUVECs. Binding and binding reinforcement values have been adjusted according to the unbinding rates: At low loading rates binding and unbinding rates are within the same order of magnitude, but one is higher depends on the binding distance. For intermediate loading rates, binding is predominant due to the reinforcement of bonds. For high load rates, bonds ultimately rupture as unbinding is predominant past the catch bond peak of maximal lifetime, estimated from [30]. Radial forces in the monolayer were reported by [62]. For protrusive forces we have selected values around four times lower than radial forces. Geometrical parameters of the model are estimated based on our experimental images, and the geometrical parameters of adhesions are extracted from [61]. Stress fiber stiffness is obtained from models of epithelial cells [29]. Membrane stiffness is within the range of values reported for two neighboring membrane ring segments in [60], and similar to measurements of cellular cortex stiffness in endothelial cells [64]. Membrane and stress fiber viscosity are within the order of magnitude of values reported for viscous drag coefficients for filament shrinkage [60]. The value of the membrane bending stiffness is within two orders of magnitude of values reported in [60] by the cell height of approximately $10 \mu m$. The medium drag coefficient is also within one order of magnitude of values used in another model for epithelial cell monolayers [29]. Typical values reported for cortical tension are of the order of $400 pN/\mu m$ [65]. If we assume that the membrane has a thickness of $20 nm$, the resulting force is around $8 pN$, which is considerably smaller than the active contraction forces. For the density of adhesion molecules, we use VE-cadherin as a proxy, given its established role as critical player to form effective cell-cell adhesions. The numerical value is estimated from

| Parameter | Symbol | Value | Source |
|---|---|---|---|
| Medium drag coefficient | $\xi$ | $4.1 \cdot 10^{-3}$ $(kg/s)$ | [29] |
| Membrane stiffness | $K_{memb}$ | $2.5 \cdot 10^{-3}$ $(kg/s^2)$ | [60] |
| Stress fiber stiffness | $K_{sf}$ | $1.25 \cdot 10^{-4}$ $(kg/s^2)$ | [29] |
| Rotational spring constant | $K_{bend}$ | $7.5 \cdot 10^{-17}$ $(Nm)$ | [60] |
| Membrane viscosity | $\eta_{memb}$ | $1.109 \cdot 10^{-3}$ $(kg/s)$ | [60] |
| Stress fiber viscosity | $\eta_{sf}$ | $1.109 \cdot 10^{-3}$ $(kg/s)$ | [60] |
| Force to normalize parameters in unbinding law | $F^0$ | 0.008 $(nN)$ | Adjusted from [30] |
| Non-dimensionalized force of catch curve in unbinding law | $\theta_c$ | 0.01 | Adjusted from [30] |
| Non-dimensionalized force of slip curve in unbinding law | $\theta_s$ | 4 | Adjusted from [30] |
| Unbinding rate coefficient for catch curve | $k_c^0$ | 0.27 $s^{-1}$ | Adjusted from [30] |
| Unbinding rate coefficient for slip curve | $k_s^0$ | 0.27 $s^{-1}$ | Adjusted from [30] |
| Binding rate for adhesions at maximum distance | $k_{on}^0$ | 15.3 $(\mu m^2/(mol \cdot s))$ | Estimated from unbinding law |
| Binding rate for adhesion reinforcement at zero force | $k_{reinf}^0$ | 11.5 $(\mu m^2/(mol \cdot s))$ | Estimated from unbinding law |
| Adhesion complex density | $\rho_{adh}$ | 21 $(mol/\mu m^2)$ | [35] |
| Limit distance for cadherin binding | $L_{bind}^{limit}$ | 0.95 $(\mu m)$ | Estimated |
| Force constant for reinforcement curve | $\lambda_{reinf}$ | 10 $nN)$ | Adjusted from unbinding law |
| Force threshold to stop applying reinforcement | $F_{reinf}^{limit}$ | 0.06 $(nN)$ | Adjusted from unbinding law |
| Adhesion complex stiffness constant per bond | $K_{adh}^0$ | $2 \cdot 10^{-4}$ $(kg/s)$ | Estimated |
| Adhesion complex equilibrium length | $L_{adh}^0$ | 0.1 $(\mu m)$ | [61] |
| Maximum number of cadherins per clutch | $n_c^{max}$ | 8 | Estimated |
| Maximum force due to radial contraction | $F_{Radial}$ | 0.775 (nN) | Adjusted from [62] |
| Maximum force due to cortical tension | $F_{cortex}$ | 0.025 (nN) | [63] |
| Maximum force due to protrusion | $F_{Prot}$ | 0.08 (nN) | Estimated |
| Force recalculation time for radial force | $t_{Radial}^{Force}$ | 25 $min$ | Estimated |
| Force recalculation time for cortical force | $t_{Cortex}^{Force}$ | 25 $min$ | Estimated |
| Force recalculation time for protrusive force | $t_{Prot}^{Force}$ | 25 $min$ | Estimated |
| Force transition time | $t_{Transition}^{Force}$ | 2 $min$ | Estimated |
| Number of nodes with similar radial force | $n_{Radial}^{Force}$ | 5 | Estimated |
| Number of nodes with similar cortical force | $n_{Cortex}^{Force}$ | 10 | Estimated |
| Number of nodes with similar protrusive force | $n_{Prot}^{Force}$ | 20 | Estimated |
| Force activation probability for radial force | $p_{Radial}^{Force}$ | 0.01 | Estimated |
| Force activation probability for cortical force | $p_{Cortex}^{Force}$ | 0.01 | Estimated |
| Force activation probability for protrusive force | $p_{Prot}^{Force}$ | 0.1 | Estimated |
| Constant for repulsion | $K_{rep}$ | $10^{-3}$ $(kg/s^2\ )$ | Estimated |
| Maximum distance to apply repulsion | $L_{rep}$ | 0.05 $(\mu m)$ | Estimated |
| Remodel rate constant | $k_{remodel}$ | $0.025 s^{-1}$ | Estimated |
| Hexagon side length | $l_{hexagon}$ | $25(\mu m)$ | Estimated |
| Distance between membrane points | $l_n$ | $625(nm)$ | Estimated |
| Minimum area for gap formation | $A_{GAP,F}$ | $2(\mu m^2)$ | Estimated |
| Area for gap closing | $A_{GAP,C}$ | $1.5(\mu m^2)$ | Estimated |
| Time step | $\Delta t$ | $1.26(s)$ | |

Table S1: Reference model parameters used in the simulation.

similar models focusing on E-cadherin in epithelial cell models [35]. The time step is a numerical parameter that is chosen sufficiently low, so that results are stable and convergent. In Fig. S15 we can see that reducing the time step does not change the results significantly. However, as we increase the time step, gap opening frequency is changing. This is because these larger time steps become of the order of magnitude of adhesion binding and unbinding time scales that affect the dynamics of the system. The choice of the time step is consequently an optimal choice that guarantees convergence of our results while being as large as possible for optimal simulation performance.

To reproduce experimental results, we performed a parameter fitting for those parameters of the model that were not fixed according to published literature values, as outlined above. Generally, our fitting was performed through adjusting published parameter values in similar systems, e.g. epithelial cells. First, we had to adjust the viscosity of the system, together with the cell generated forces and the stiffness of both stress fibers and membrane segments. These parameters combined, control the overall velocity at which the nodes move. This is important to reproduce the velocity at which cells deform and move, and to fit the experimentally observed lifetime of the gaps. Binding and unbinding parameters are very important to reproduce gap opening frequencies, since they control the rupture and binding of the adhesion complexes. The binding law is estimated and we set the values so they are consistent with the unbinding law rate from literature values. Finally, bending stiffness has a strong influence on the location where the gaps are generated (border vs vertex), and was consequently adjusted so that computational results matched our experimental data.

We varied selected parameters within certain ranges in the results section of our main text. We focused on fold changes that have maximal impact on the shown results. Data for physiological ranges that can motivate these fold changes are rarely available. We considered fold changes typically within one order of magnitude from the reference case, which is likely obtainable through experiments or in different physiological conditions. However, even if such fold changes are not physiologically relevant, they demonstrate the principle impact of the represented physical parameters, and therefore, their importance, on gap formation.

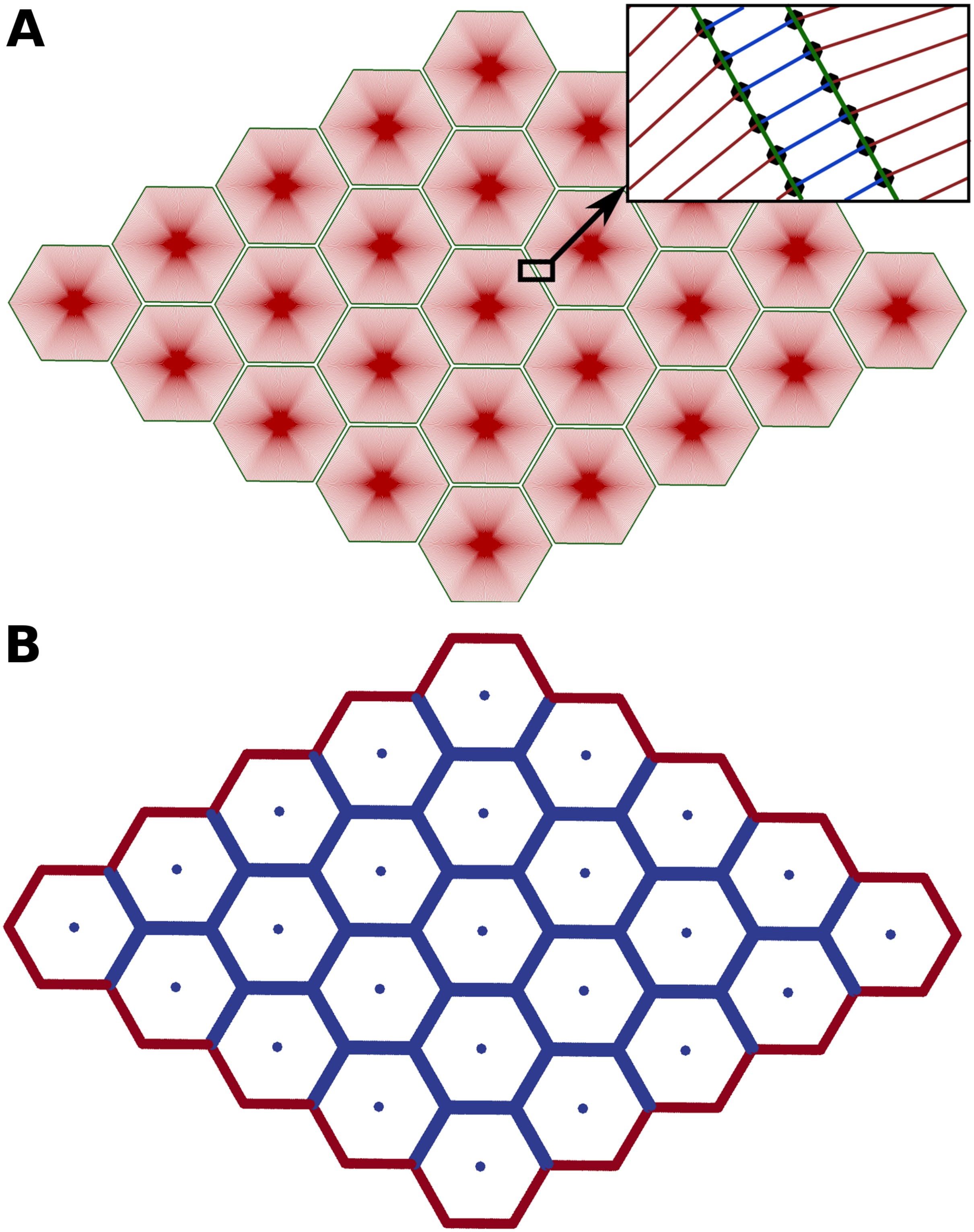


Figure S5: **Model of the endothelial monolayer.** (A) Cells with a hexagonal shape are in a rest state and fully bound to their neighboring cells. Cell membrane (green), stress fibers (red), cadherin complexes (blue), membrane points (black). (B) Boundary conditions: Points in the boundary of the monolayer (red) are fixed. In blue are membrane points and the cell centers.

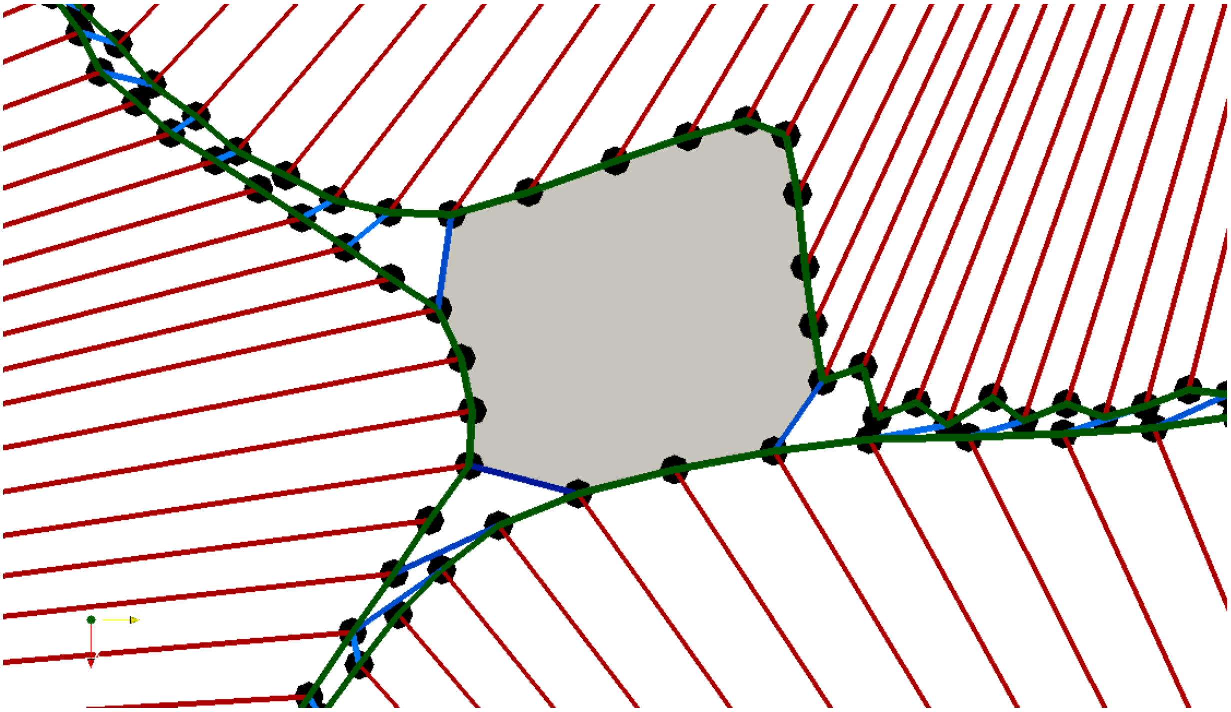

Figure S6: **Paracellular gap.** A gap (grey area) is delimited by the cell membrane (green) and the adhesion bonds binding the cells (blue). Red: cell stress fibers. Black dots: Membrane points

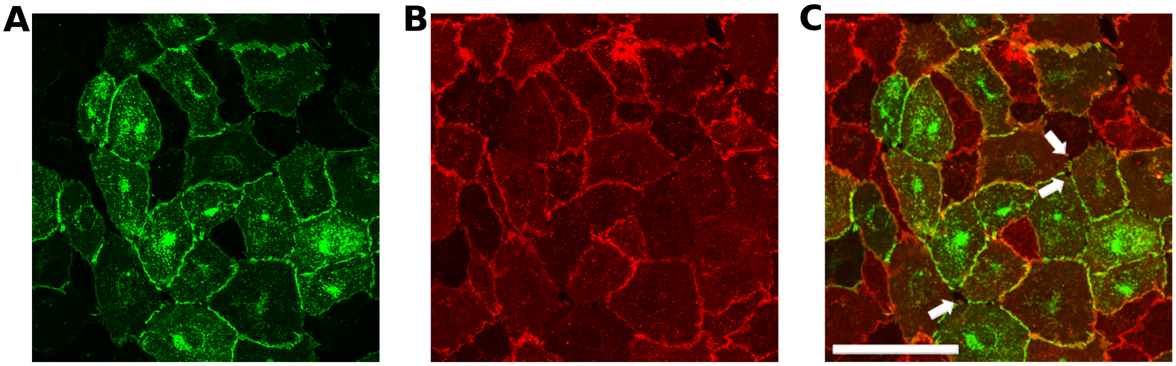


Figure S7: **Gaps in VE-cadherin correspond to gaps in CD31.** Endothelial monolayer stained with VE-cadherin (green, A) and CD31 (red, B). C: Merged image confirms that gaps observed within the VE-cadherin mediated cell-cell adhesions are also present within CD31, indicating that gaps seen in VE-cadherin are real physical gaps between the cells. Scale bar $100\mu m$.

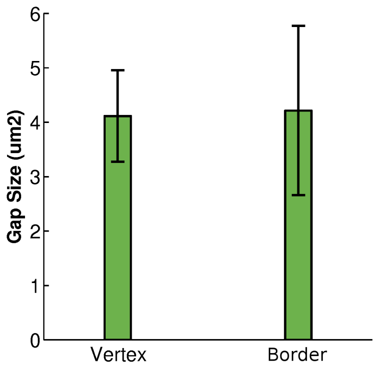


Figure S8: **Gap sizes predicted from simulations with reference parameters.** Average size of the gaps generated at the vertices and borders. Parameters are the reference values as in Table S1 and error bars correspond to standard deviation of sample=30.

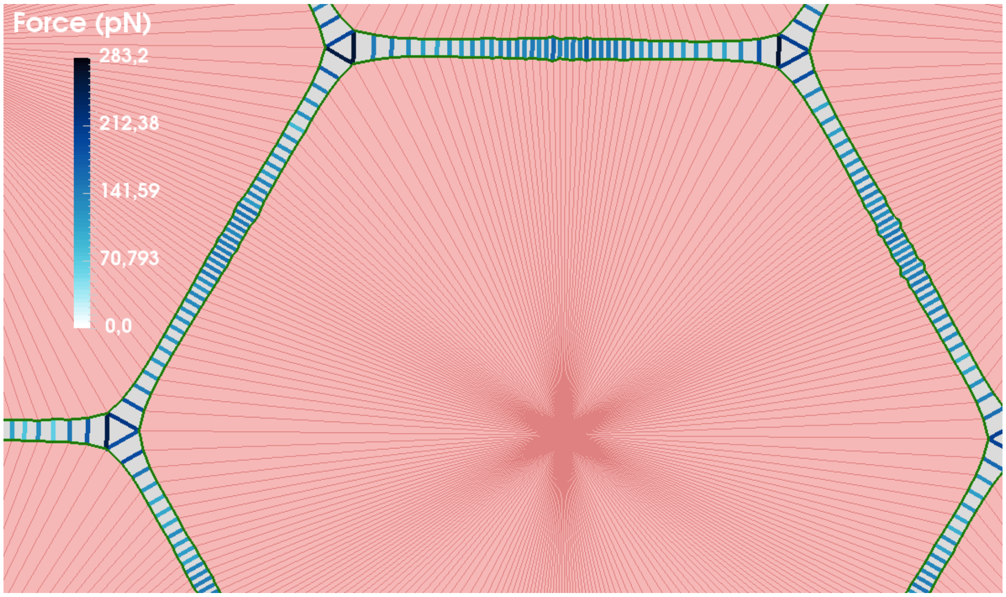


Figure S9: **Stresses on the cell-cell adhesions.** Homogeneous contractions are applied to all the hexagonal cells in the monolayer. Stresses concentrate on the adhesions at vertices, as opposed to the adhesions at border.

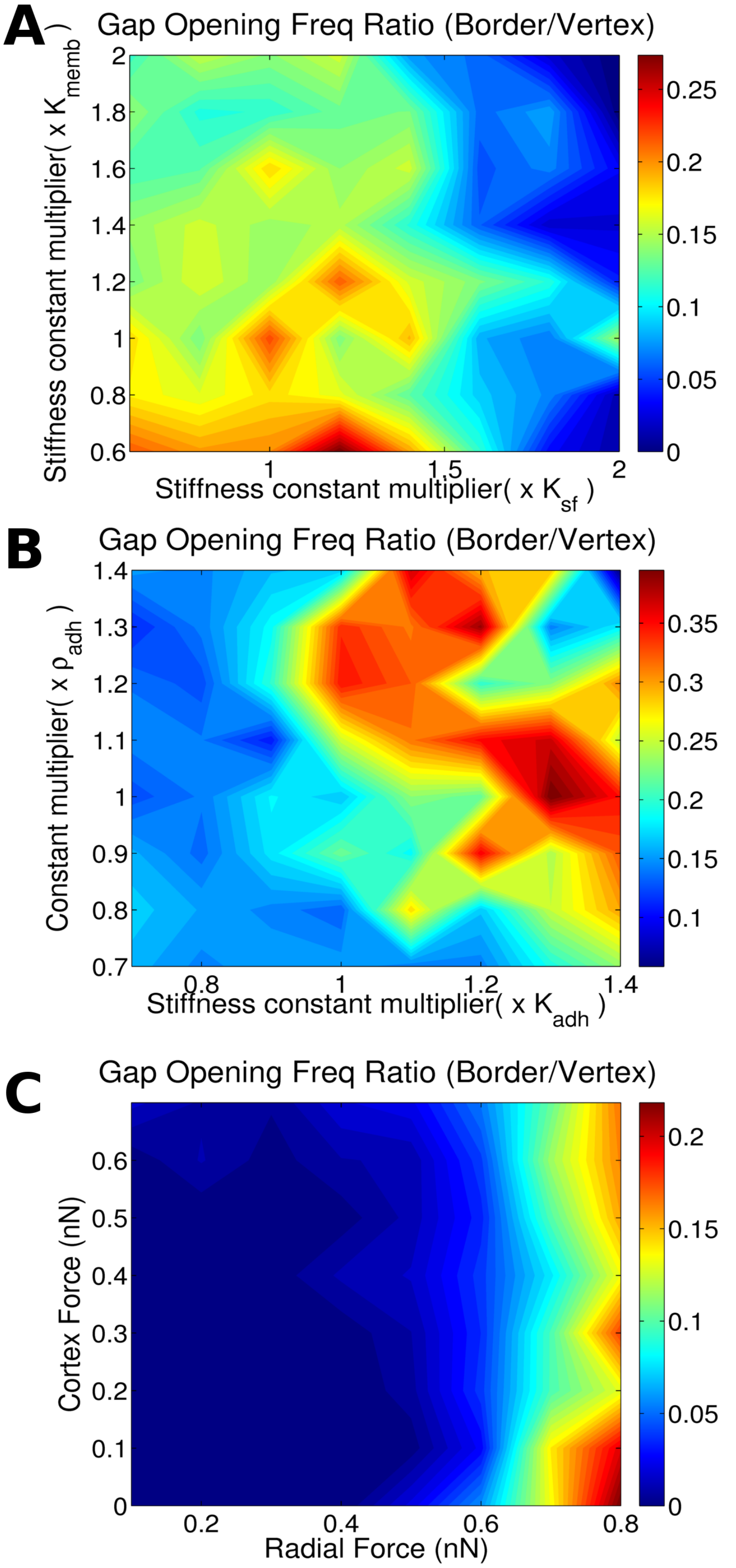


Figure S10: **Effect of two parameter variation on gap opening location.** Shown is the ratio of gaps that occur at a two cell border divided by the gaps that originate at a three cell vertex. A shows results varying membrane and stress fiber stiffness. B shows properties of cell-cell junction are changed: cadherin stiffness versus cadherin density (binding rate). C shows results for varying cortical and radial force.

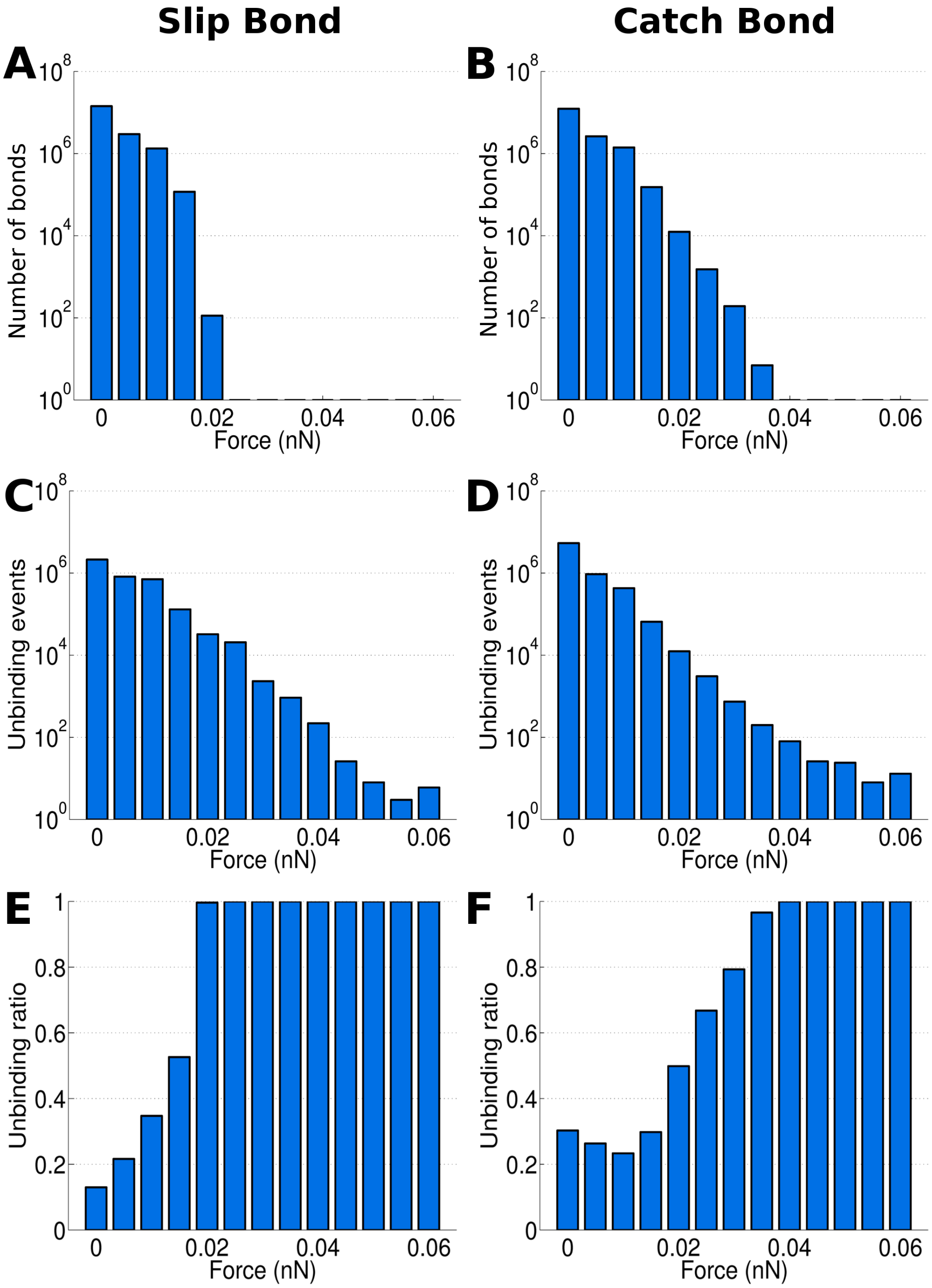


Figure S11: **Forces on bonds, comparing a pure slip bond and the catch bond law used as reference in the paper.** First row shows force histogram of cadherins that are bound. Second row cadherins force at which cadherins unbind. Third row shows the ratio obtained by dividing unbound cadheins by the sum of unbound cadherins and bound cadherins ($ub/(ub + b)$, where ub and b corresponds to unbound and bound cadherins respectively.)

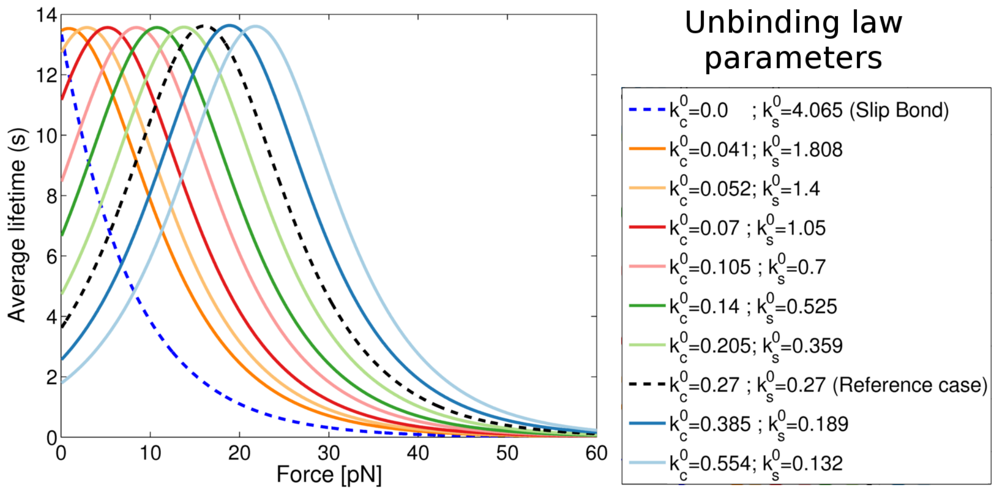


Figure S12: **Shift of force of maximal catch bond lifetime.** Lifetime average for the bond in dependence on the force for different unbinding laws. Legend shows the parameter variation to obtain the different curves.

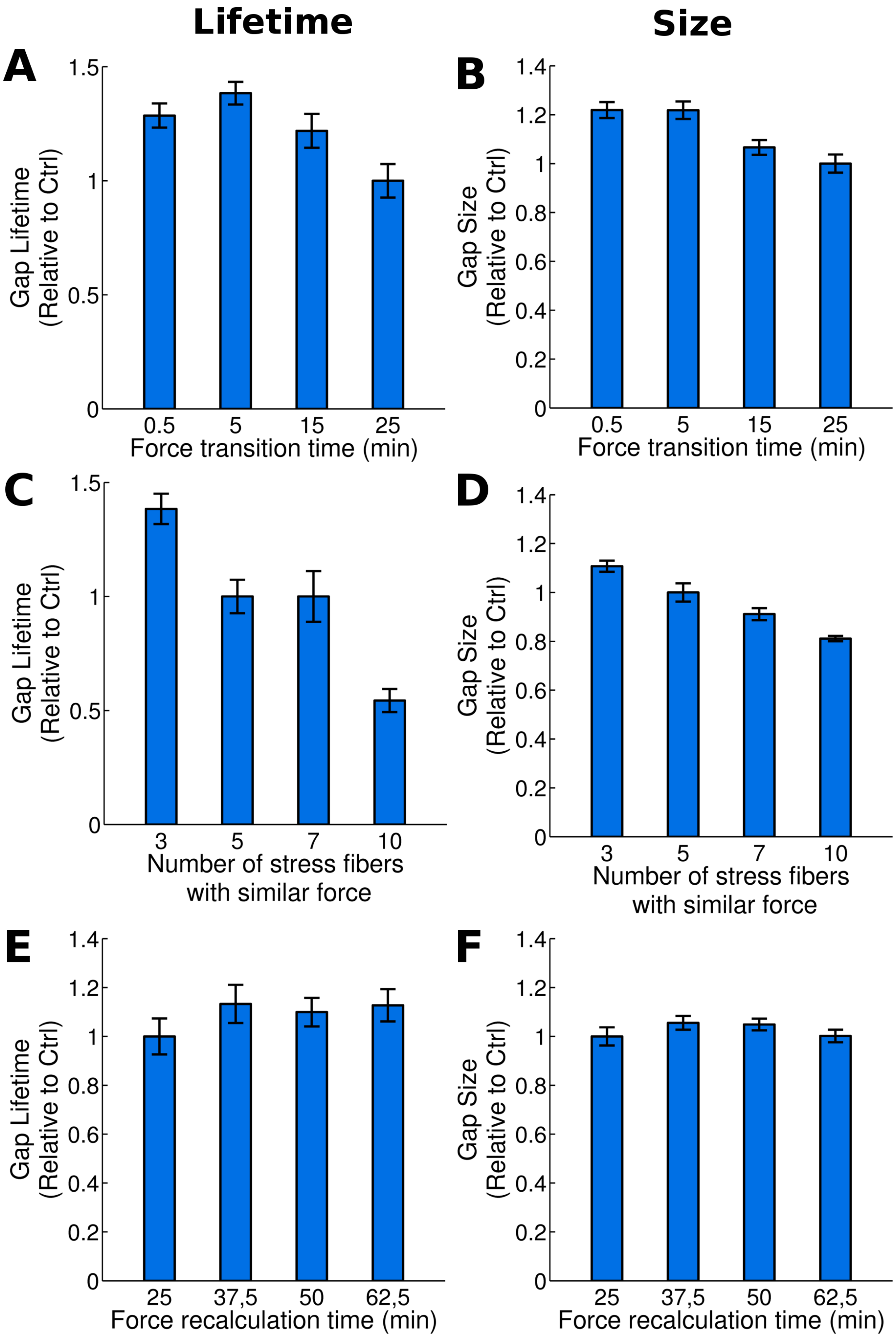


Figure S13: **Effect of force application on gap size and lifetime.** Corresponds to Fig. 4. Left column corresponds to lifetime and right column to size. (A, B): Changes in the transition time of the application of the recalculated forces. Longer time means smoother force changes. (C, D) Variation in the number of stress fibers over which the same force is distributed. (E, F) Variation in force fluctuation time for all types of forces considered in the model. Error bars show to the standard error.

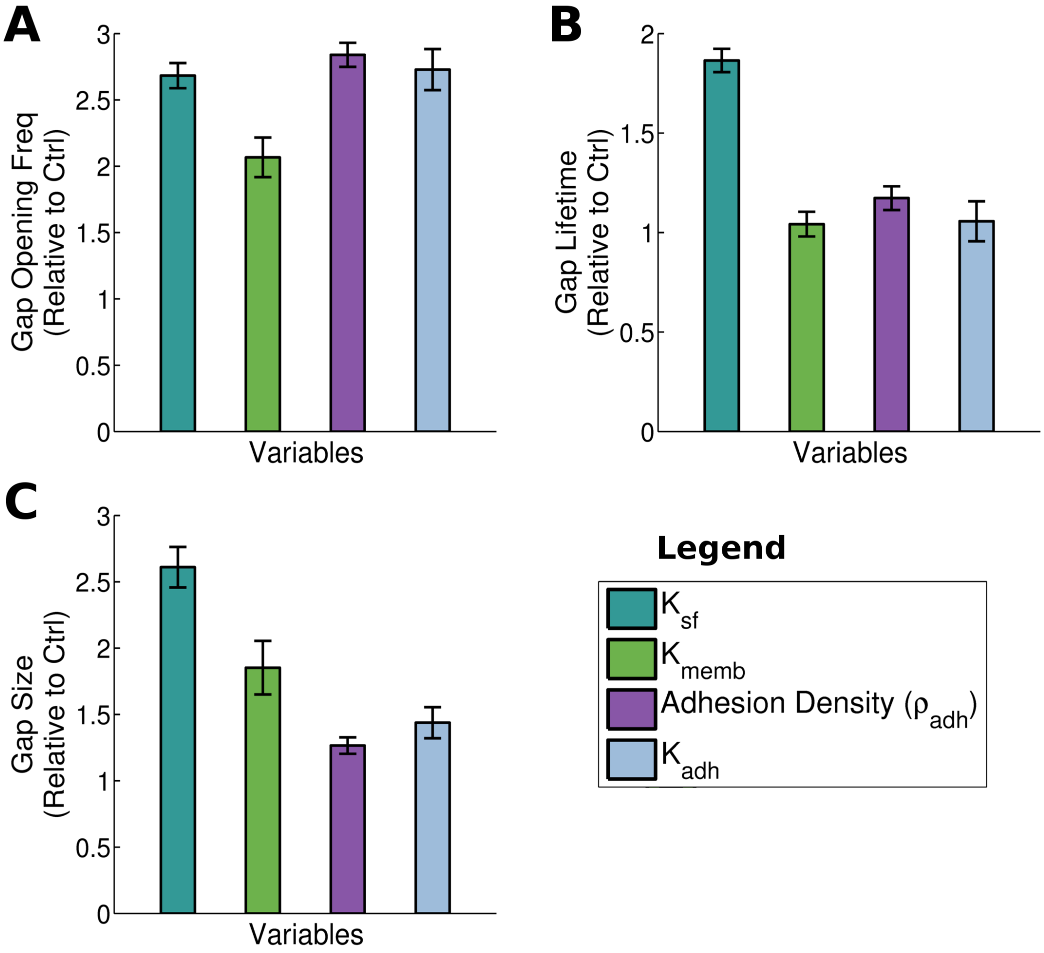


Figure S14: **Interplay of adhesion and cell mechanical properties controls different aspects of gap opening dynamics.** Error bars show to the standard error. All parameters have been reduced one order of magnitude ($x10^{-1}$). (A) Gap opening frequency. (B) Average lifetime of the gaps. (C) Average size of the gaps.

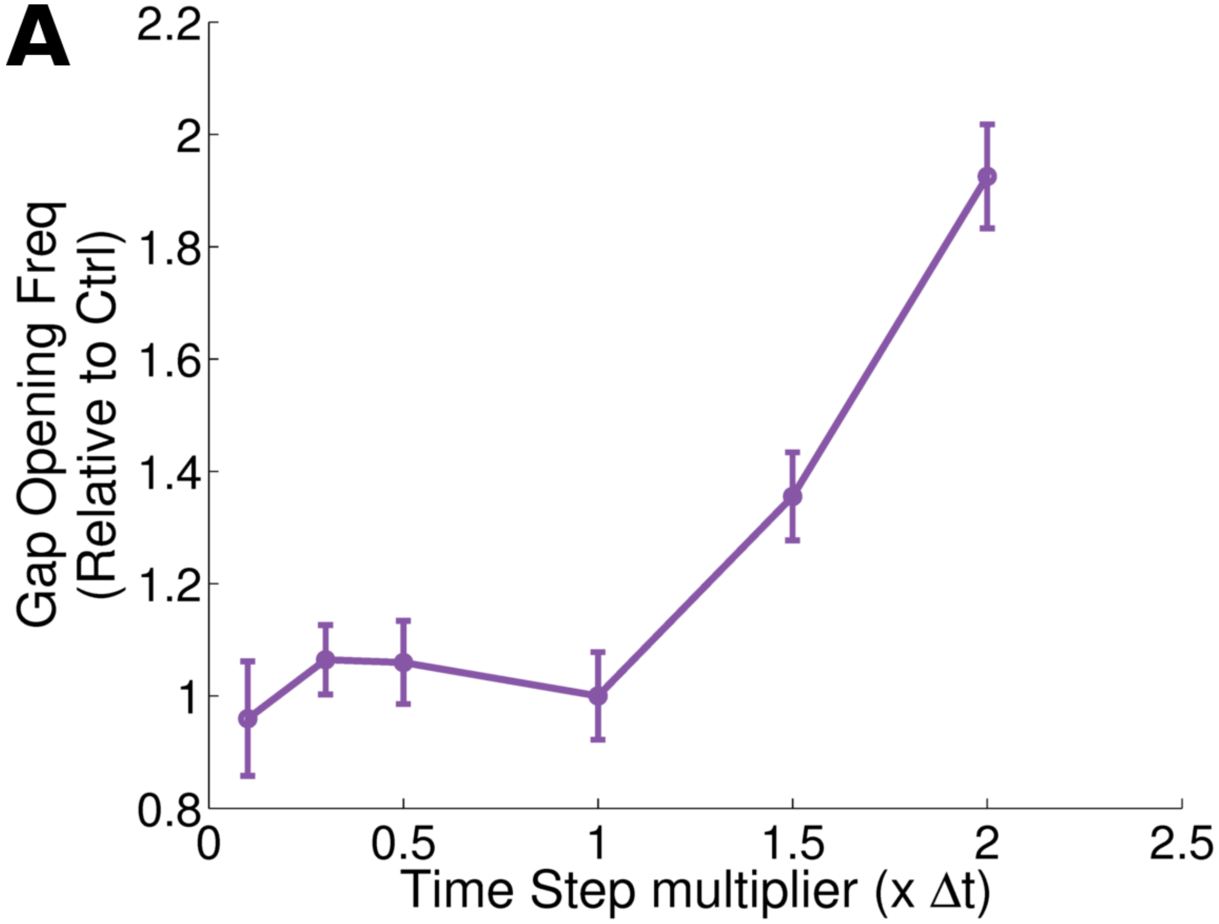


Figure S15: **Time step analysis.** The gap opening frequency depends on the time step used in our numerical simulations. Note that the time step multiplier is relative to the reference case (multiplier = 1). Error bars are the standard error. The results confirm that the time step selected for the reference case is low enough to ensure convergence of the results.

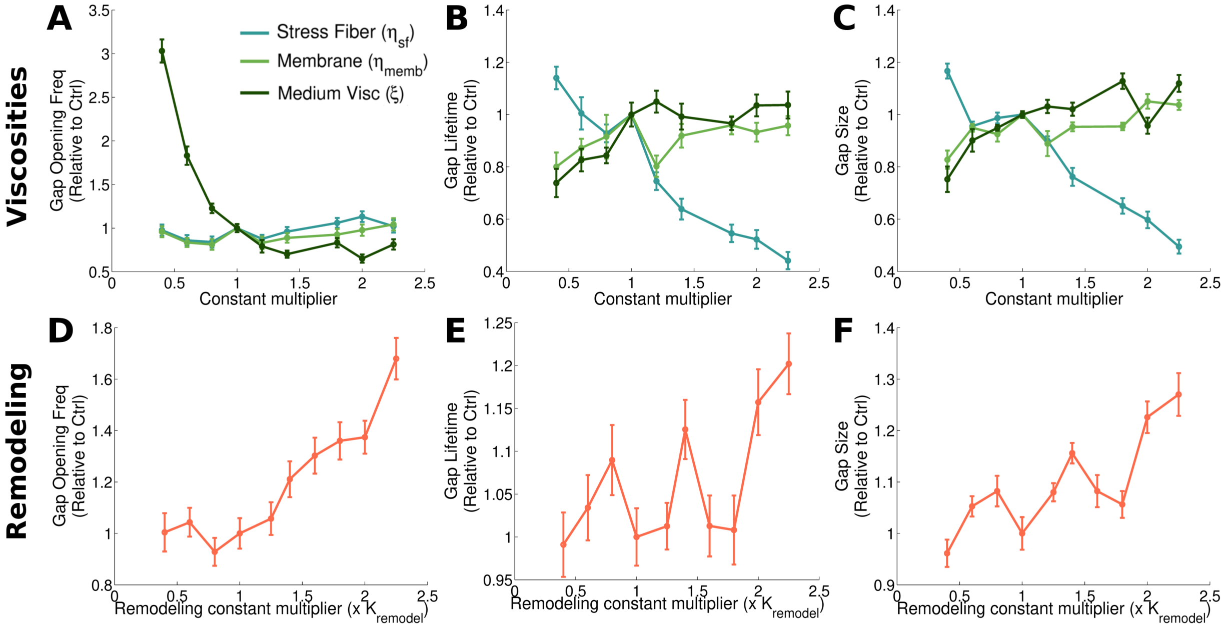


Figure S16: **Effect of the viscosities and the remodeling rate on the gap opening dynamics.** Gap opening frequency, average lifetime and size in each column. Note that the point where x and y coordinates are 1 corresponds to the reference case. Error bars represent standard error. In the first row (A, B, C), results for medium and dashpot viscosities of the stress fibers and membrane and varied. Increasing viscosity reduces node movement, stabilizing monolayer dynamics. Medium viscosity has a higher effect on gap opening dynamics since it affects the overall timescale of all mechanical parts of the model. The stress fiber dashpot strongly influences gap lifetime and size; this is similar to the dominating effect of stress fiber stiffness over membrane stiffness on gap lifetime and size (Fig. 3A,B) . The second row (D, E, F) shows the effect of varying the constant for remodeling rate. Increasing the remodeling rate implies that cells are able to adapt their permanent shapes faster in response to deformations. Therefore, the frequency of gap openings increases with the remodeling rate (D). The gap lifetime and size broadly also increase, but less strongly then the opening frequency.

Movie S1: **Simulation of the endothelial monolayer dynamics.** Gaps are more likely to appear in the vertex of three cells than at a two cell border. Green denotes the cell membrane, red the inside of a cell, with darker red being the stress fibers. Parameters are the reference values as in Table S1

Movie S2: **Experimental observation of an endothelial monolayer dynamics.** Dynamics of a monolayer of HUVEC cells, corresponding to Fig. 1D-F

Movie S3: **Stresses on the cell-cell adhesions.** Homogeneous contractions are applied to a hexagonal cell, showing that stresses naturally concentrate on the adhesions at vertices, as opposed to the adhesions at the border. This leads to a faster gap generation at these areas.

Movie S4: **Altered monolayer dynamics due to low stiffness in the stress fibers.** Not only gap opening frequency is increased under these conditions but also, gaps are critically larger compared to the reference case.

Movie S5: **Altered monolayer dynamics due to high stiffness in the stress fibers.** Gap opening frequency is strongly suppressed for very stiff stress fibers.

Movie S6: **Altered monolayer dynamics due to low bending stiffness.** Membranes can easily deform when forces are applied, reducing gap formation.

Movie S7: **Altered monolayer dynamics due to high bending stiffness.** Cells tend to be more rounded, provoking a concentration of stress at the adhesions at the vertices and leading to gap generation in these zones. Gaps are bigger and difficult to close.

Movie S8: **Altered monolayer dynamics due to slip bonds.** Gap opening frequency is clearly increased under these conditions compared to the reference case (based on catch bonds).

Movie S9: **Cancer cell extravasation occurring at endothelial cell vertex.** MDA-MB-231 tdTomato (red) extravasating through a HUVEC endothelial monolayer at a vertex. Endothelial junctions are visualized via VE-cadherin GFP (green). After successful transmigration, cancer cells spreads and migrates below the monolayer, followed by the re-sealing of the endothelial gap. Images are taken every 12 minutes.
